# Class II LBD genes *ZmLBD5* and *ZmLBD33* regulate gibberellin and abscisic acid biosynthesis

**DOI:** 10.1101/2021.04.08.439062

**Authors:** Jing Xiong, Xuanjun Feng, Weixiao Zhang, Xianqiu Wang, Yue Hu, Xuemei Zhang, Fengkai Wu, Wei Guo, Wubing Xie, Qingjun Wang, Jie Xu, Yanli Lu

**Author notes:** These authors contribute equally to this work (Jing Xiong, Xuanjun Feng). Senior author (Xuanjun Feng). Author for contact: Yanli Lu. The author responsible for distribution of materials integral to the findings presented in this article in accordance with the policy described in the Instructions for Authors (www.plantphysiol.org) is: Yanli Lu. **Author Contributions** X.F. and Y.L. designed the research. J.X., W.Z., X.W., Y.H. and X.Z. performed the main part of experiment. X.F., Q.W, W.G., W.X. and F.W. performed the field investigation of plant phenotype. J.X. and Y.L. are responsible for managing materials related to the project. X.F. and J.X. analyzed data and prepared the figures. X.F., J.X. and Y.L. wrote the manuscript.

## Abstract

Lateral organ boundaries domain (LBD) proteins are plant-specific transcription factors. Class I *LBD* members are widely reported to be pivotal for organ development, however, the role of class II members is unknown in cereal crops. Class II LBD proteins are distinguished from class I by the lack of a Gly-Ala-Ser (GAS) peptide and leucine-zipper-like coiled-coil domain, which is thought to be essential for protein dimerization. In this study, ZmLBD5 and ZmLBD33 form homo- and hetero-dimers, like class I members. At seedling stage, *ZmLBD5* promoted biomass accumulation (shoot dry weight and root dry weight), root development (root length, root number, and root volume), and organ expansion (leaf area), while *ZmLBD33* repressed these processes and display a dwarf phenotype. Both *ZmLBD5* and *ZmLBD33* displayed negative roles in drought tolerance mainly by increasing stomatal density and stomatal aperture. RNA sequencing, gene ontology enrichment analysis, and transient luciferase expression assays indicated that *ZmLBD5* and *ZmLBD33* are mainly involved in the regulation of the *TPS*-*KS*-*GA2ox* gene module, which comprises key enzymatic genes upstream of GA and ABA biosynthesis. GA_1_ content increased in *ZmLBD5-*overexpressing seedlings, while GA_3_ and abscisic acid content decreased in both transgenic seedlings. Consequently, exogenous GA_1_ or GA_3_ undoubtedly rescued the dwarf phenotype of *ZmLBD33*-overexpressing plants, with GA_1_ performing better. The study of *ZmLBD5* and *ZmLBD33* sheds light on the function of the class II *LBD* gene family in maize.

## INTRODUCTION

Gibberellins (GAs) and abscisic acid (ABA) are two phytohormones derived from isopentenyl diphosphate (IPP). In plastids, IPP is converted to monoterpenes (C_10_) and geranylgeranyl diphosphate (C_20_, GGPP). At GGPP the pathway in plastids branches out in several directions, with separate pathways leading to the *ent*-kaurenoids and GAs (C_20_), the phytyl side-chain of chlorophyll (C_20_), phytoene and carotenoids (C_40_), and the nonaprenyl (C_45_) side-chain of plastoquinone (Hedden and Sponsel, 2015; He et al., 2020). Thus, manipulation of one of the pathways can have significant effects on the flux through other branches, and any regulation in the pathway upstream of GGPP will affect all branches. In GA biosynthesis, GGPP is first cyclized to *ent*-copalyl diphosphate (CPP) by CPP synthases (CPSs) and then converted to *ent*-kaurene by *ent*-kaurene synthases (KSs) (Hedden and Sponsel, 2015; He et al., 2020). This is followed by oxidative reactions catalyzed by cytochrome P450 oxygenases and 2-oxoglutarate-dependent oxygenases that regulate the dynamic balance between activation and deactivation of GAs (Hedden and Proebsting, 1999; Fu et al., 2016).

GAs and ABA are two of the most studied phytohormones in plants. The essential role of GAs in shoot elongation has been demonstrated clearly in a large number of studies (Spray et al., 1996; Teng et al., 2013; Chen et al., 2014; Nagai et al., 2020). In addition, GAs regulate various aspects of plants, such as flowering (Bao et al., 2020), root development (Zimmermann et al., 2010), seed development (White et al., 2000), and stress response (Colebrook et al., 2014), and the “green revolution” of farming occurred largely owing to the application of *GA20ox* knockout crops (Hedden, 2003). ABA has multiple functions, and it is well known to ameliorate abiotic stress. Under the stress of drought, ABA accumulates rapidly and plays a positive role in drought tolerance by regulating multiple processes at different tiers, such as the expression of ABA-responsive genes, stomatal closure, root growth, and the production of protective metabolites (Mehrotra et al., 2014). ABA is mainly produced from the cleavage of carotenoids (Sponsel and Hedden, 2010; Xiong and Zhu, 2003), which are co-derived from GGPP along with GAs (Sponsel and Hedden, 2010). Thus, the crosstalk between GAs and ABA biosynthesis is universal.

Lateral organ boundaries domain (LBD) proteins are plant-specific transcription factors (Shuai et al., 2002; Majer and Hochholdinger, 2011; Xu et al., 2016). Characteristically, they comprise a cysteine C-block (CX2CX6CX3C) required for DNA-binding activity, a Gly-Ala-Ser (GAS) block, and a leucine zipper-like coiled-coil motif (LX6LX3LX6L) responsible for protein dimerization (Shuai et al., 2002; Majer and Hochholdinger, 2011; Xu et al., 2016). Based on the conserved domains, LBD genes have been identified and classified into two groups (class I and class II) (Shuai et al., 2002; Majer and Hochholdinger, 2011; Zhang et al., 2014; Yu et al., 2020). Class I, comprising most of the members, is characterized by a GAS and leucine-zipper-like coiled-coil domain, while class II members have no or an incomplete GAS and leucine zipper-like coiled-coil domain. It is thought that these differences underlie the functional diversity between class I and class II members (Majer and Hochholdinger, 2011; Xu et al., 2016). Class I members have been mostly reported to play an important role in auxin response and plant development (Majer and Hochholdinger, 2011), such as maintaining the indeterminate cell state in the shoot apical meristem (SAM) (Semiarti et al., 2001; Iwakawa et al., 2007), female gametophyte development (Evans, 2007), inflorescence architecture (Bortiri et al., 2006), and formation of seminal and shoot-borne root primordia (Taramino et al., 2007; Majer et al., 2012; Xu et al., 2015). In contrast, there are few reports about class II members, and their functions are unclear. The class II *LBD* genes that have been characterized thus far are not involved in development but in metabolism, such as anthocyanin biosynthesis and nitrogen metabolism (Rubin et al., 2009; Albinsky et al., 2010; Majer and Hochholdinger, 2011). Although the phylogenetic characteristics of the *LBD* family in maize have been reported (Zhang et al., 2014), there are no reports about the function of class II *LBD*s in cereal crops.

In this study, two class II *LBD* maize genes, *LBD5* and *LBD33*, with high identity (Zhang et al., 2014), were investigated. Here we found homo- and hetero-dimerization in both LBD5 and LBD33. This is at odds with the hypothesis that the GAS and leucine-zipper-like coiled-coil domain is essential for dimerization in LBDs. In addition, LBD5 and LBD33 play a role in the regulation of the core module in GA biosynthesis and affect the final output of GAs and ABA. This work may contribute to the initial understanding of the biological function of cereal class II *LBD* genes in cereal crops, and provide a new insight into the response of drought in maize.

## RESULTS

### LBD5 and LBD33 can form homodimers and heterodimers

Given that the GAS and leucine-zipper-like coiled-coil domain of class I members is essential for protein dimerization and class II members are characterized by the lack of or an incomplete domain, the ability of LBD5 and LBD33 to dimerize was tested. Although LBD5 and LBD33 have no GAS and leucine-zipper-like coiled-coil domain, homo- and hetero-dimerization were clearly detected through Y2H and BiFC assays (Fig. 1, A and B). Additionally, LBD5 and LBD33 displayed weak interactions with the class I member LBD44 (Fig. 1, A and B). To identify the critical domain for dimerization, LBD5 and LBD33 were cleaved optionally at two sites, and six kinds of peptides were obtained for Y2H assays (Supplemental Fig. S1A). Parts A, B, and C represent the N terminal C-block (CX2CX6CX3C), the GAS and leucine-zipper-like coiled-coil (LX6LX3LX6L) domain, and the C terminal domain, respectively (Fig. S1A). Unfortunately, we failed to clearly identify which part is responsible for protein-protein interaction (Supplemental Fig. S1B), but we proposed that the C part may have a positive effect on dimerization both in LBD5 and LBD33 (Supplemental Fig. S1B).

**Figure 1.**
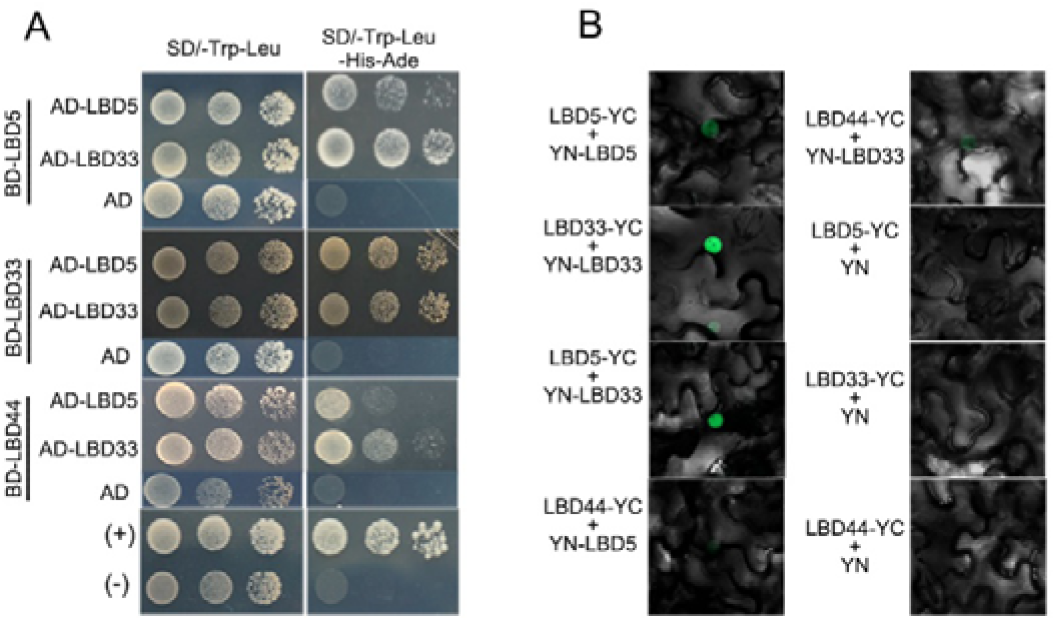
Protein-protein interaction analyzed by yeast two-hybrid and bimolecular fluorescence complementation. Fll codon region of *ZmLBD5, ZmLBD33*, and *ZmLBD44* were used here. (A) The yeast cells harboring the indicated plastid combinations were grown on nonselective (SD/-Trp/-Leu) or selective (SD/-Trp/-Leu/-His/-Ade) medium. Cells were diluted in three concentrations from left to right. (B) ZmLBD5, ZmLBD33 and ZmLBD44 was individually cloned into pXYc104 and pXYn106 vectors, and fused with C-terminal (YC) or N-terminal (YN) of YFP. The indicated plastid combinations were transiently co-expressed in tobacco.

### Effects of LBD5 and LBD33 on organ development and drought tolerance

To investigate the role of *LBD5* and *LBD33* in plant development or environmental stimuli, first their tissue-specific expression and stimuli responses were profiled. *LBD5* and *LBD33* were widely expressed in various tissues and relatively higher in roots (Supplemental Fig. S2A). Therefore, the response of *LBD5* and *LBD33* to eight phytohormones related to stress response and organ development was investigated in the roots of seedlings. *LBD5* and *LBD33* were induced most strongly by GA_3_ and JA, and these stimuli had similar effects (Supplemental Fig. S2B). In addition, responses to PEG, nitrogen deficiency, and phosphorus deficiency were investigated in the root and leaf. *LBD5* and *LBD33* were remarkably induced by PEG-mimic drought stress and nitrogen deficiency in the root. However, responses in the leaf were weak and differed slightly between *LBD5* and *LBD33* (Supplemental Fig. S1,C and D).

LBD5 and LBD33 were overexpressed, which were called LBD5(OE) and LBD33(OE) hereafter, in a wild-type maize line KN5585. More than seven transgenic lines of each gene were constructed and transcript levels of *LBD5* and *LBD33* were measured. Three representative transgenic lines of each gene were used for further study after the detection of expression levels (Supplemental Fig. S3A). Homozygous lines were hybridized with the wild type, and the hybrid F1 progenies were used for phenotype investigation. During germination and the early seedling stage there was no difference between LBD5(OE), LBD33(OE), and the wild type (Supplemental Fig. S3B). About 6 days after germination, an obvious dwarf phenotype of LBD33(OE) was observed. About 12 days after germination, LBD5(OE) was noticeably taller than the wild type. Detailed analysis showed that the shoot length, leaf area, and biomass were remarkably affected by the over expression of *LBD5* and *LBD33*, and all of these phenotypic values increased in LBD5(OE) but decreased in LBD33(OE) (Fig. 3 and Supplemental Fig. S3D). To investigate the change at the micro-level, leaf cell length and width was observed. As expected, cells were longer in LBD5(OE) and shorter in LBD33(OE) than that in the wild type (Supplemental Fig. S4). Unexpectedly, LBD33(OE) plants possessed narrower leaves but wider cells and LBD5(OE) possessed wider leaves but narrower cells than the wild type (Supplemental Fig. S4). We thus speculated that the leaf cell number may be lower in LBD33(OE) and higher in LBD5(OE). To investigate the effects on below-ground parts, seedlings were grown in hydroponic conditions. The root number, primary root length, root surface area, root volume, shoot fresh weight, and root fresh weight were all greater in LBD5(OE) and lesser in LBD33(OE) than that in the wild type (Supplemental Fig. S3,C and D). However, the difference in root dry weight between LBD5(OE), LBD33(OE), and the wild type was not significant when grown in soil (Fig. 2B).

**Figure 2.**
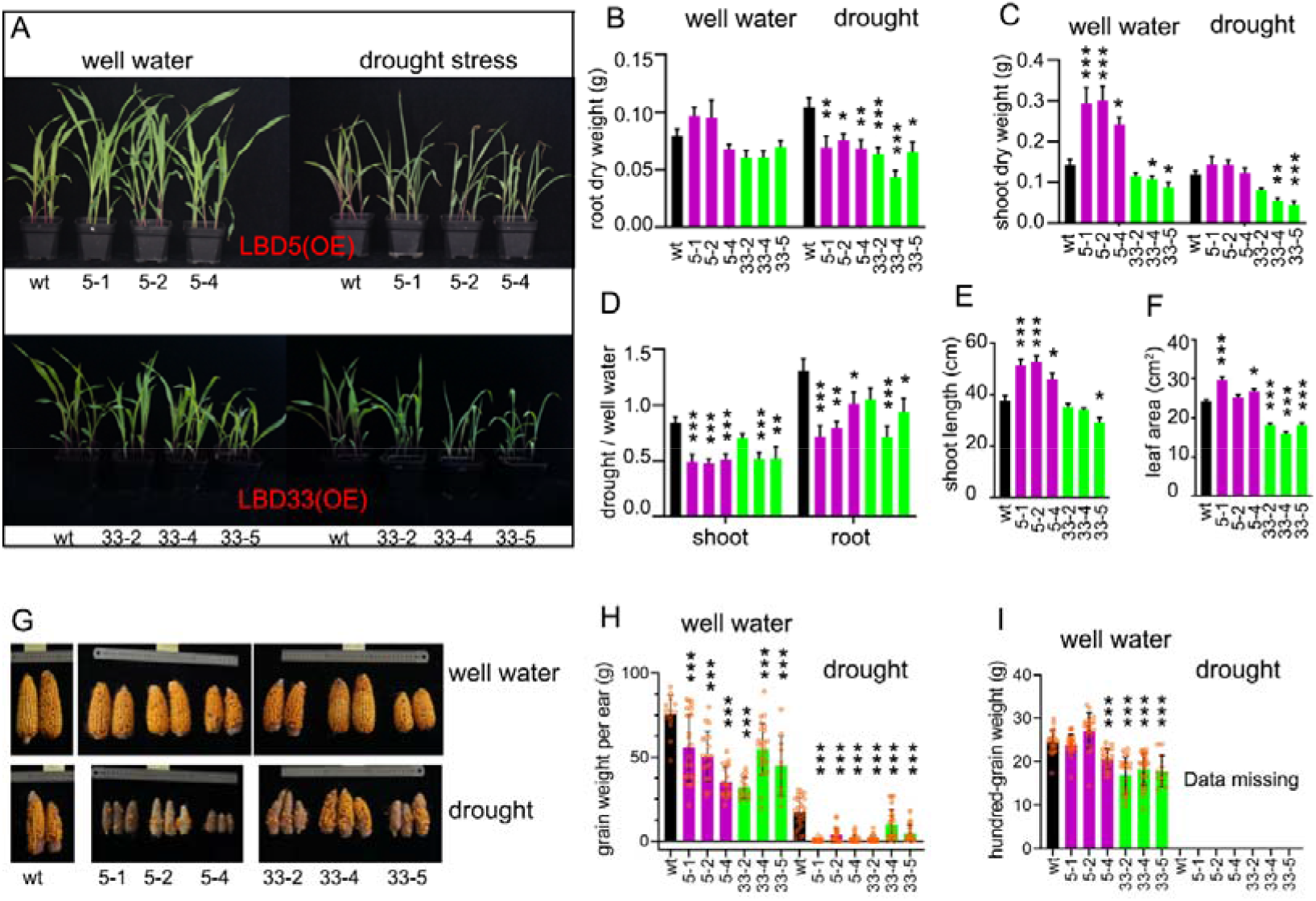
*ZmLBD5* or *ZmLBD33* over-expressed plants are drought sensitive. (A) Wild type and individual transgenic lines were grown in separate pot exposed to well water or drought stress conditions. Quantitative description the phenotypes of root dry weight (B), shoot dry weight (C), ratio of dry weight between drought stress and well water conditions (D), shoot length (E) and leaf area (F). Ear phenotype (G), grain weight per ear (H), and hundred-grain weight (I) of transgenic plants and wild type in field upon well water and drought conditions. Asterisks on bar represent the difference compared with wild type is significant.

**Figure 3.**
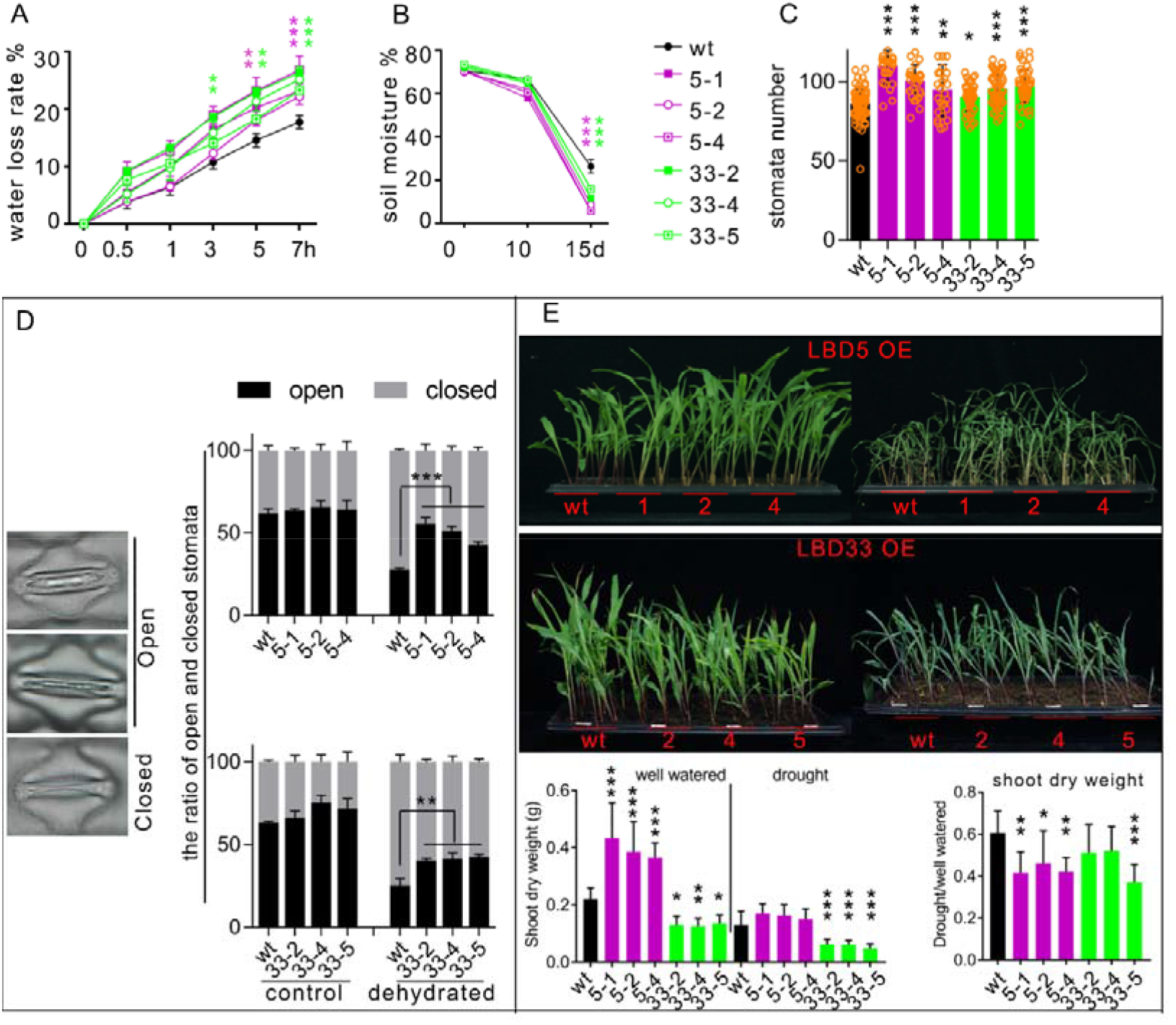
Overexpression of *ZmLBD5* or *ZmLBD33* promotes water loss. (A) Water loss rate of detached leaf. (B) Soil moisture of each pot grown with wild type or different transgenic plants after the outage of water. (C) Stomata number of the third leaf on abaxial surface. (D) The ratio of open and closed stomata 15 minutes after detachment. Represented open and closed stomata were shown in the left panel. (E) Phenotype of seedlings before and after drought stress when different lines of transgenic plants and wild type were grown in the same pot. In panels (A), (B) and (D), if all three lines were significant different with the wild type asterisk was labeled.

Given the strong response of *LBD5* and *LBD33* to PEG-mimic drought stress, wild type and different transgenic lines were grown in separate pots and exposed to water deficiency. Both LBD5(OE) and LBD33(OE) displayed earlier withering than the wild type, and this effect was more remarkable in LBD5(OE) than in LBD33(OE) (Fig. 2A). The biomass penalty under water deficient conditions also indicated that LBD5(OE) and LBD33(OE) were more sensitive to drought stress than the wild type when grown in separate pots (Fig. 2, B-D).

However, after anthesis in the field, the ear height and the internode length of lines 5-2 and 5-4 were shorter than that of the wild type (Supplemental Fig. S5C and Table S1). However, the effect of LBD5 on plant height was not clear, because line 5-1 was taller, line 5-2 was shorter, and line 5-4 was similar when compared with the wild type (Supplemental Fig. S5,A and B). The increase in plant height of line 5-1 may be attribute to the larger number of internodes (Supplemental Fig. S5D). Thus, we speculate that LBD5 may have a positive effect on internode number and a negative effect on internode length. LBD33(OE) plants were significantly shorter than the wild type owing to the decrease in internode length but not internode number (Supplemental Fig. S5, A-D and Table S1). Grain yields upon well-watering and drought stress conditions supported the drought sensitive phenotype of LBD5(OE) and LBD33(OE) (Fig. 2, G-I). In addition, *LBD5* and *LBD33* negatively regulate grain yield with different mechanism. *LBD33* decreased ear size and hundred-grain weight simultaneously, while *LBD5* mainly decreased ear size (Fig. 2, G-I and Supplemental Fig. S5,E and F.

### LBD5 and LBD33 promote water loss by increasing stomatal density and stomatal aperture

To unravel the underlying mechanism by which LBD5(OE) and LBD33(OE) increased sensitivity to drought stress, the rates of water loss (RWL) of detached leaves were measured. LBD5(OE) and LBD33(OE) had significantly higher RWL than the wild type (Fig. 3A). The stronger water loss in LBD5(OE) and LBD33(OE) may have, consequently, resulted in earlier soil drought after the outage of water. Thus, the soil moisture content (SMC) was analyzed. Fifteen days after the outage of water SMC was lowest in LBD5(OE) and remarkably lower in LBD33 (OE) than in the wild type (Fig. 3B). Further, the stomatal density and open-closed-ratio (OCR) were investigated. To measure OCR, leaves were detached from 12-day-old seedlings and dehydrated for 12 minutes in original growing conditions. Interestingly, LBD5(OE) and LBD33(OE) had higher stomatal density and OCR than the wild type (Fig. 3,C and D and Supplemental Fig. S6A).

In addition, stomatal aperture after ABA or H_2_O_2_ stimulation was investigated. Detached leaves were pretreated with stomatal opening buffer to get the maximum degree of stomatal openness. Then, ABA or H_2_O_2_ was added, and stomatal aperture was fixed at different times. The stomatal aperture of LBD5(OE) and LBD33(OE) was greater than that of wild type, upon both ABA and H_2_O_2_ treatment (Supplemental Fig. S6,B and C). ABA and H_2_O_2_ are both powerful stimulators of stomatal closure. These results indicated that overexpression of *LBD5* or *LBD33* could reduce the sensitivity of stomata to ABA and H_2_O_2_.

To determine if there were other factors that made LBD5(OE) and LBD33(OE) more sensitive to drought stress, transgenic and wild-type plants were grown in the same pots to test the difference in drought tolerance. LBD5(OE), LBD33(OE), and the wild type withered at almost the same time after the outage of water. However, the penalty of shoot dry weight upon drought stress was greater in LBD5(OE) and LBD33(OE) (Fig. 3E). These results suggest that enhanced water loss from stomata was one cause of drought sensitivity in LBD5(OE) and LBD33(OE) plants.

### LBD5 and LBD33 may function in the GGPP–CPP–kaurene/acid–GA metabolic pathway

As transcription factors, overexpression of LBD5 or LBD33 may widely affect the expression of downstream target genes, leading to the final phenotype. Therefore, RNA-seq was performed using 12-day-old seedlings to profile the change of gene expression in LBD5(OE) and LBD33(OE) compared with the wild type. A total of 1844 and 1174 differentially expressed genes (DEGs) were identified in LBD5 (OE) and LBD33(OE), respectively (Fig. 4A and Supplemental Table S2). There were 666 members in the intersection of DEGs between LBD5(OE) and LBD33(OE), and almost all of these genes displayed a similar expression change in LBD5(OE) and LBD33(OE) when compared to the wild type (Fig. 4A). Of the 666 shared DEGs, 417 were down regulated (Fig. 4A). When DEGs from LBD5(OE) and LBD33(OE) were combined, the 3018 total genes could be clustered into 6 groups based on the change in expression level (Fig. 4B). Almost all of the shared 666 genes were found in group 2 or group 5, in which genes displayed a similar expression change in LBD5(OE) and LBD33(OE) compared to the wild type. It is plausible that the similar phenotype of LBD5(OE) and LBD33(OE) is caused by the 666 shared DEGs or the other members of group 2 and group 5.

**Figure 4.**
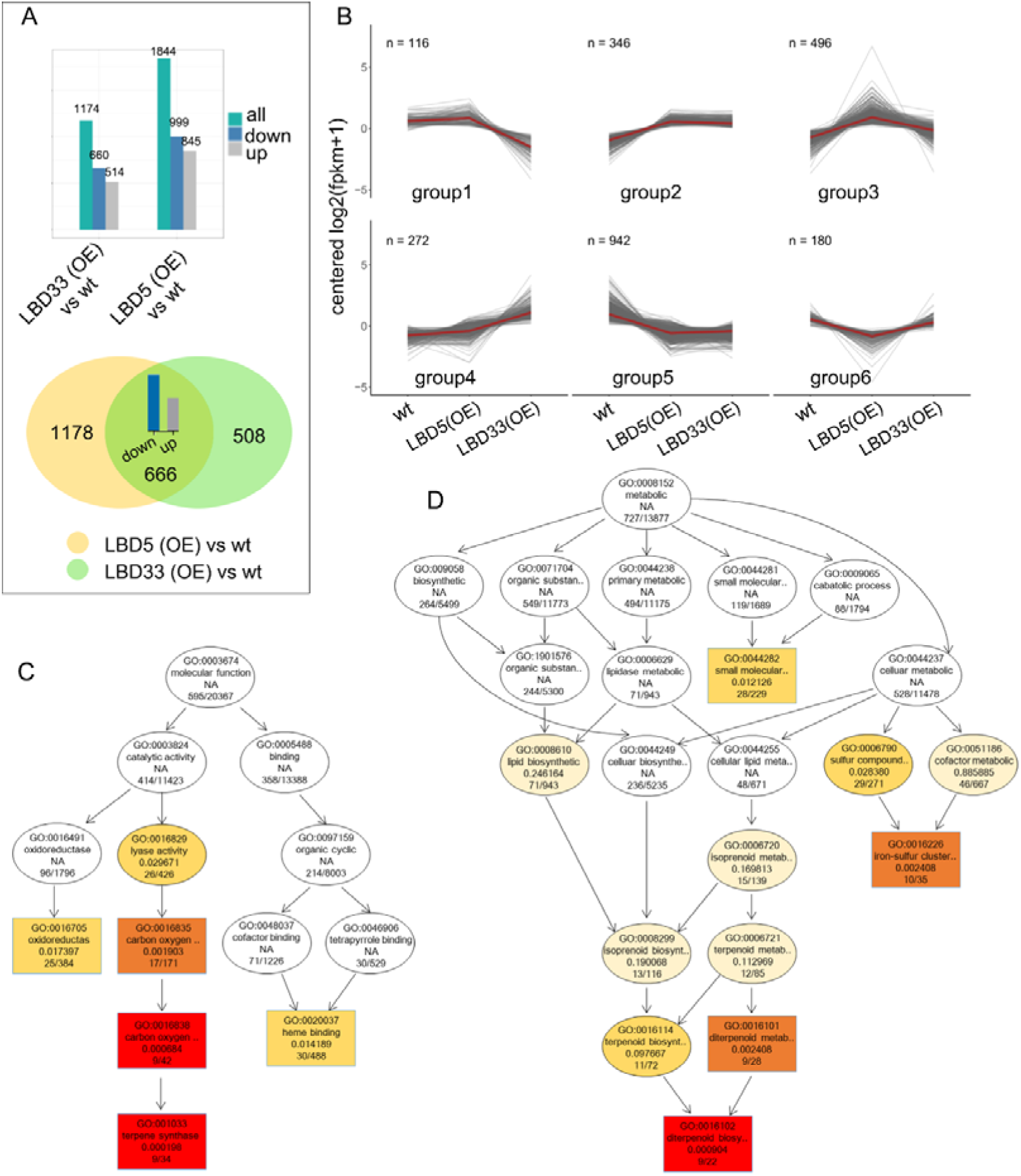
RNA-seq indicates *ZmLBD5* and *ZmLBD33* function in terpenoid metabolic pathway. (A) Different expressed gene (DEG) number in *LBD5* or *LBD33* overexpressed plants compared with the wild type. The fold change larger than 2 or smaller than 0.5 were determined as DEGs. (B) All of the DEGs were classified into 6 groups by the centered and logarithmic transformed expression levels in wild type, LBD5 (OE) and LBD33 (OE). Directed Acyclic Graph of gene ontology enrichment analysis by using DEGs from *LBD5* overexpressed plants (C) and *LBD33* overexpressed plants (D). The four lines of information in each circle represent GO number, function annotation, P-value of GO enrichment significance test, and DEG number/background gene number, respectively. The darker the circle, the higher the enrichment.

Further, DEGs of LBD5(OE) and LBD33(OE) were used for Gene Ontology (GO) enrichment analysis. Results provided a similar and reliable biologic process for LBD5(OE) and LBD33(OE), namely terpenoid biosynthesis (Fig. 4C and Supplemental Table S3). A total of 15 DEGs annotated in terpenoid biosynthesis were mainly involved in the GGPP–CPP–kaurene/acid–GA pathway (Fig. 5A). Five of the six genes involved in GGPP biosynthesis were down-regulated in LBD5(OE) and LBD33(OE) (Fig. 5A). We speculated that those 15 genes may be downstream targets of LBD5 and LBD33.

**Figure 5.**
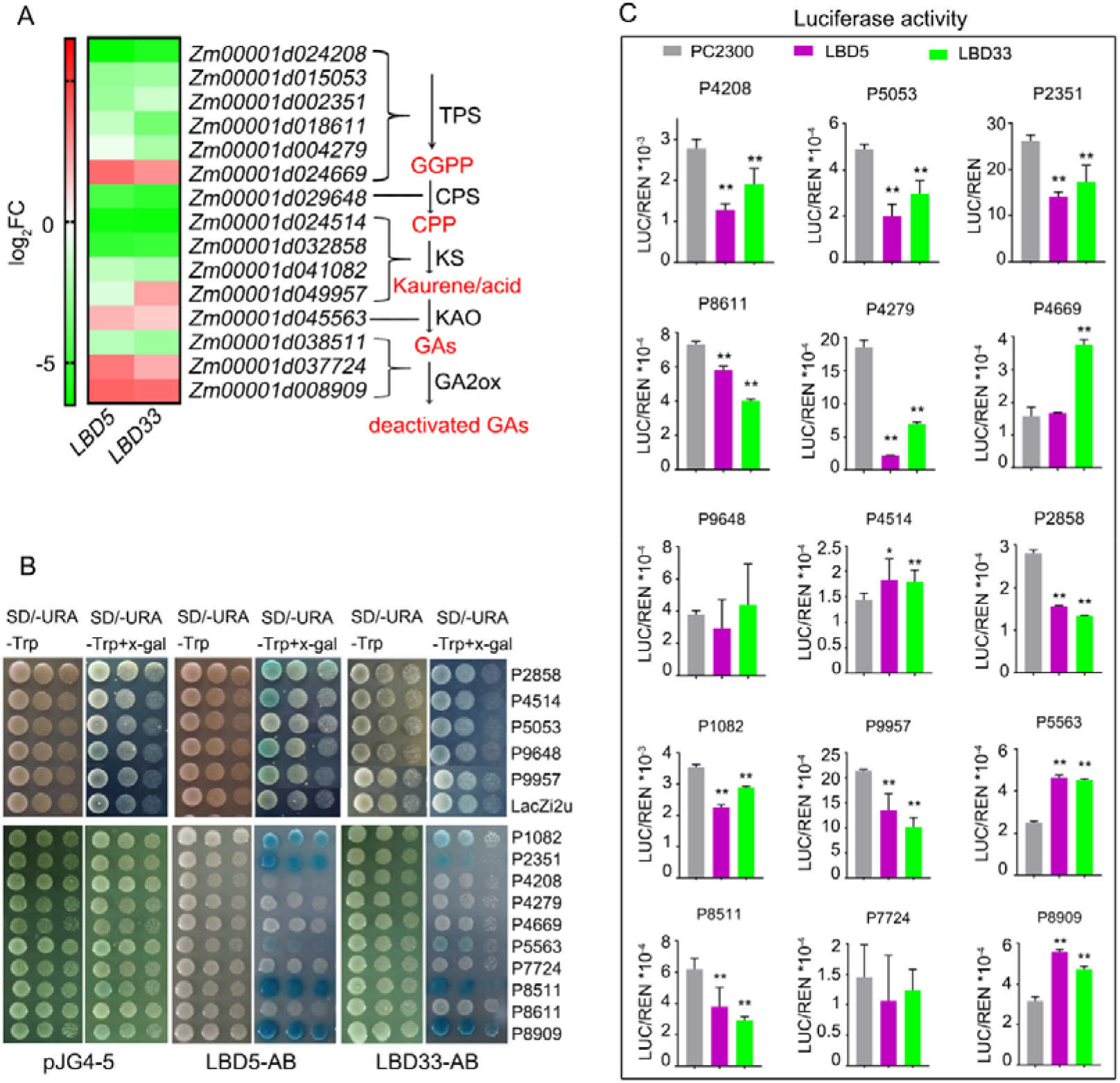
ZmLBD5 and ZmLBD33 directly regulate *TPS-KS-GA2ox* gene module. (A) 15 DEGs in the most enriched GO terms in figure 4C and 4D were listed. The expression fold change (FC) was converted to a logarithmic scale with base 2 and shown as different color. TPS: terpene synthase; CPS: *ent*-copalyl diphosphate synthase; KS: *ent*-kaurene synthase; KAO: *ent*-kaurenoic acid oxidase; GA2ox: GA 2-oxidase; GGPP: geranylgeranyl diphosphate; CPP: *ent*-copalyl diphosphate. (B) Y1H testing whether AB-segment of LBD5 and LBD33 bind to the promoters of five candidate genes. The yeast cells harboring the indicated plastid combinations were grown on nonselective (SD/-Ura/-Trp) or color development (SD/-Ura/-Trp/+x-gal) medium. The last four number of the candidate gene name was used to represent corresponding gene. Cells were diluted in three concentrations from left to right. (C) dual-luciferase reporter system was used to investigate the regulation of LBD5/LBD33 on candidate genes. Asterisks on bar represent the difference compared with wild type is significant.

### LBD5 and LBD33 directly regulate the *TPS*-*KS*-*GA2ox* gene module

To investigate whether LBD5 and LBD33 directly regulate the transcription of candidate target genes, promoters of the candidate genes were cloned for yeast one-hybrid assays. Firstly, the full length of LBD5 and LBD33 were used for the Y1H assays along with 5 candidate promoters. However, none of the five promoter-driven *LacZ* reporter genes could be activated by LBD5 or LBD33 (Supplemental Fig. S7A). Given that most genes were down-regulated by LBD5 and LBD33, we speculated that full length LBD5 and LBD33 may have transcriptional inhibitory activity and block the activation of reporter gene. Therefore, LBD5 and LBD33 were cleaved optionally at two sites and six kinds of peptides were obtained, as shown in Figure S1A, to test the transcriptional inhibitory activity. Full length LBD5, LBD5-BC, LBD33-B, and LBD33-BC displayed remarkable transcriptional inhibitory activity (Supplemental Fig. S7B). LBD5-AB and LBD33-AB, which contained the C-block for DNA binding, had no transcriptional inhibitory activity. Therefore, LBD5-AB and LBD33-AB were then used for Y1H assays along with 15 promoters. LBD5-AB and LBD33-AB could bind to 9 and 5 promoters, respectively (Fig. 5B).

Given that LBD proteins are plant specific transcription factors, the regulation of 15 candidate target genes by LBD5 and LBD33 was investigated in plant cells. The promoters of candidate genes were cloned to drive the expression of the *LUC* reporter gene. CaMV35S-controlled *LBD5* or *LBD33* was fused with *GFP* and transiently co-expressed with each candidate-promoter-driven *LUC* in tobacco. Compared with the fold-change of transcription level, luciferase enzymatic activity assays showed that 13 and 11 candidate promoters displayed similar behavior upon the overexpression of *LBD5* and *LBD33*, respectively (Fig. 5,A and B). Taking Y1H and dual-luciferase expression assays into consideration, we identified that LBD5 and LBD33 directly regulated 7 and 5 of the 15 candidate target genes, respectively.

### LBD5 and LBD33 affect seedling size by modulating GA biosynthesis

In plants, ABA is mainly produced from the cleavage of carotenoids (Sponsel and Hedden, 2010; Xiong, 2003), which are co-derived from GGPP along with GAs. Aforementioned results indicated that LBD5 and LBD33 negatively regulate drought tolerance by inhibiting stomatal closure, a process in which ABA is important. Therefore, concentrations of ABA and GAs (the main bioactive form GA_1_, GA_3_, GA_4_, and GA_7_) (Hedden and Phillips, 2000), two kinds of phytohormone downstream of GGPP, were measured using 12-day-old seedlings. ABA and GA_3_ contents were lower in both LBD5(OE) and LBD33(OE) than in the wild type (Fig. 6A). Interestingly, GA_1_ content was higher in LBD5(OE) than in the wild type (Fig. 6A). This may explain why LBD5(OE) seedlings were taller than the wild type. However, it is difficult to determine the causal gene from the 15 candidate genes based on the subtly different responses to LBD5 and LBD33 overexpression. Most redox reactions in GA metabolism are catalyzed by cytochrome P450s (He et al., 2020; Hedden and Sponsel, 2015; Sponsel and Hedden, 2010), and 44 cytochrome P450 members were identified in the top enriched GO term (Supplemental Table S3). Thus, the expression change of these 44 members may also contribute to the change in GAs and ABA in LBD5(OE) and LBD33(OE).

**Figure 6.**
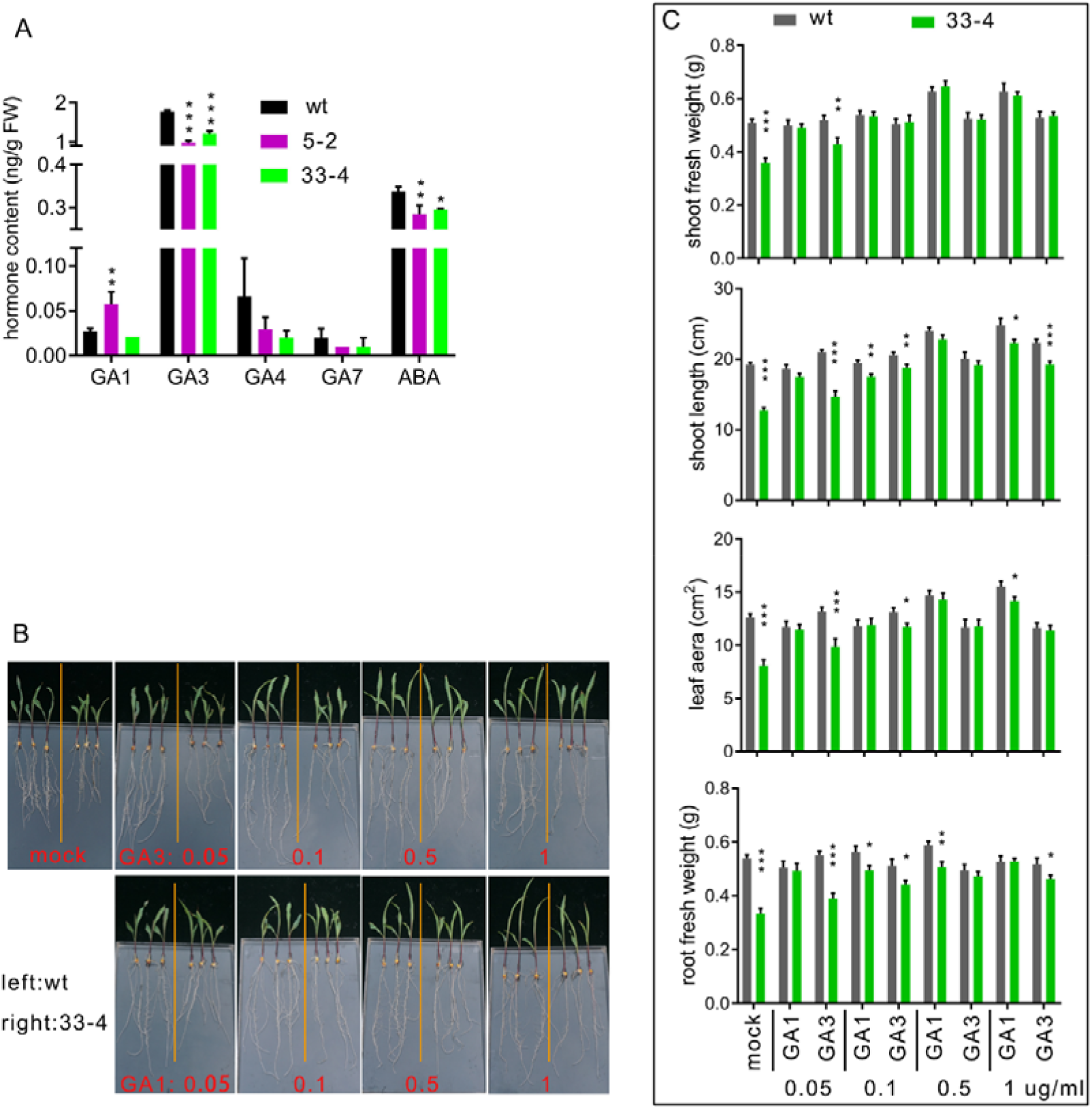
Exogenous GA_1_ and GA_3_ rescue the dwarf phenotype of *ZmLBD33* overexpressed plants. (A) Endogenous content of GAs and ABA of 12-days-old seedlings. (B) Phenotype and (C) quantitative description of shoot length, leaf area, shoot fresh weight and root fresh weight. Asterisks on bar represent the difference compared with wild type is significant.

GAs widely affect plant development, particularly stem elongation (Spray et al., 1996; Teng et al., 2013; Chen et al., 2014; Nagai et al., 2020). To test the causality of GA_1_ and GA_3_ content on the dwarf phenotype of LBD33(OE) seedlings, exogenous GA_1_ and GA_3_ were applied in hydroponic conditions. The application of GA_1_ and GA_3_ remarkably promoted the growth of LBD33(OE) seedlings and reduced the difference between LBD33(OE) and the wild type (Fig. 6A). GA_1_ displayed a well phenotypic compensatory effect at 0.05 µg/mL (Fig. 6,B and C). However, the effect of GA_3_ was weak at 0.05 µg/mL, and a higher concentration was needed to better rescue the dwarf phenotype of LBD33(OE) seedlings (Fig. 6,B and C). Therefore, GA_1_ may exhibit the dominant effect on shoot growth and elongation at the seedling stage in maize.

## DISCUSSION

Class I members of the LBD gene family are involved in the regulation of almost all aspects of plant development, including embryo, root, leaf, and inflorescence development (Majer and Hochholdinger, 2011). However, there are few reports to date on the function of class II members. Characterized class II members, *AtLBD37, AtLBD38*, and *AtLBD39*, are not involved in development but in anthocyanin biosynthesis and nitrogen metabolism (Scheible et al., 2004; Rubin et al., 2009; Albinsky et al., 2010; Majer and Hochholdinger, 2011). Here, we investigated the function of two members of the class II *LBD* genes in maize. Multiple lines of evidence suggested that *ZmLBD5* and *ZmLBD33* are involved in the regulation of terpenoid metabolism, and consequently determine GA and ABA content, thus affecting plant development and drought response.

Class I members of the LBD protein family are distinguished from class II members by the existence of the GAS and leucine-zipper-like coiled-coil domain (Majer and Hochholdinger, 2011; Xu et al., 2016). This domain is thought to be required for protein dimerization (Majer and Hochholdinger, 2011; Xu et al., 2016). In this study, LBD5 and LBD33, which are class II members lacking a typical GAS and leucine-zipper-like coiled-coil domain, formed homodimers and heterodimers like class I members, implying the typical GAS and leucine-zipper-like coiled-coil domain is not essential for dimerization. However, no domain responsible for dimerization has been identified in LBD5 and LBD33. Heterodimerization between LBD5 and LBD33 was strong. However, the interaction was weak when LBD44, a class I member, was used to form heterodimers with LBD5 or LBD33. These results imply that similarity between sequences may facilitate dimerization between LBD monomers.

DEGs in *LBD5-* and *LBD33-*overexpressing plants compared to the wild type are perfectly enriched in the same biological process upstream of GA biosynthesis, which was reminiscent of the similar and drastic responses of *LBD5* and *LBD33* to GA_3_ treatment. GA_3_ content decreased in both LBD5(OE) and LBD33(OE), which is consistent with the down-regulation of most DEGs in the GGPP–CPP–kaurene/acid–GA metabolic pathway. In line with the down-regulation of most DEGs, LBD5 and LBD33 have transcriptional inhibitory activity. Application of exogenous GA_1_ and GA_3_ clearly restored the dwarf phenotype of LBD33(OE), particularly for shoot length and shoot fresh weight, indicating that decreased GA_1_ and GA_3_ content are the immediate cause of the dwarf phenotype. Although the expression change of most enriched genes was similar, GA_1_ increased in LBD5(OE) and remained unchanged in LBD33(OE), which may explain the taller phenotype of LBD5(OE). GA_1_ has been identified in 86 plants, more than any other GA, and studies utilizing single gene dwarf mutants have shown that it is the major bioactive form involved in stem elongation in *Zea mays* and *Pisum sativum* (Sponsel and Hedden, 2010; Spray et al., 1996). Our results also attest that GA_1_ has a higher bioactivity, because low concentrations of GA_1_ rescued the dwarf phenotype of LBD33(OE) as well as high concentrations of GA_3_.

ABA is the most important signal in drought response. Under water deficient conditions, ABA accumulates rapidly and leads to stomatal closure to limit transpirational water loss (Mehrotra et al., 2014). In *LBD5*- and *LBD33*-overexpressing plants, the ABA content was decreased probably due to the down-regulation of genes involved in the biosynthesis of the upstream GGPP. As a result, LBD5(OE) and LBD33(OE) had larger stomatal apertures and enhanced transpirational water loss compared with the wild type. However, the mechanism by which *LBD5* and *LBD33* reduce the sensitivity of stomata to exogenous ABA and H_2_O_2_ is not clear.

In addition, *LBD5* and *LBD33* increased stomatal density. Although there are few reports on the involvement of *LBD* genes in the regulation of stomatal density, GID1, a very important receptor of GA, plays a negative role in stomatal density in rice (Du et al., 2015). The *gid1* mutant displays extreme dwarfism, increased stomatal density and decreased stomatal sensitivity to water deficiency (Du et al., 2015). Here, the phenotypes of LBD5(OE) and LBD33(OE) were similar to that of *gid1*. Further studies are needed to determine if the decrease of GA_3_ in LBD5(OE) and LBD33(OE) plants mimics *gid1* or if there exists underlying causality.

In conclusion, *ZmLBD5* and *ZmLBD33* are mainly involved in the regulation of the *TPS*-*KS*-*GA2ox* gene module, which comprises key enzymatic genes upstream of GA and ABA biosynthesis. Subtle differences in the regulation of these genes led to GA_1_ accumulation and a taller phenotype in LBD5(OE) seedlings. Overexpression of *LBD5* and *LBD33* inhibited GA_3_ and ABA biosynthesis and resulted in a dwarf phenotype in LBD33(OE) and drought sensitive phenotype in both LBD5(OE) and LBD33(OE). The study of *ZmLBD5* and *ZmLBD33* sheds light on the function of the class II *LBD* gene family in maize.

## MATERIALS AND METHODS

### Plant growth and phenotyping

To analyze the expression patterns of LBD5 and LBD33 in different maize tissues, the primary roots and coleoptiles were collected from seedlings germinated for 3 days (VE stage); seminal roots and leaves were collected at the three-leaf stage (V1 stage); and aerial root, stem, ear leaf, husk, silk, immature ears, and tassels were collected from the V13 stage in a maize inbred line (B73) for RNA isolation.

For stress and phytohormone treatment, two-leaf-stage seedlings were transferred to Hoagland nutrient solution in a greenhouse with a 14-h light/10-h dark photoperiod at 28°C and grown to the three-leaf-stage. Then, seedlings were subjected to polyethylene glycol 6000 (PEG6000) (20% w/v), nitrogen deficiency (0.1 mM nitrogen), phosphorus deficiency (without phosphorus), ABA (10 μM), GA_3_ (1 μM), ethephon (50 μM), indoleacetic acid (IAA) (5 nM), jasmonic acid (JA) (20 μM), salicylic acid (SA) (2 mM), 6-benzyl aminopurine (6-BA) (4 µM), or brassinolide (100 nM). For PEG, nitrogen deficiency, and phosphorus deficiency, root and leaf tissues were harvested after 0, 1, 3, 6, 12, 24, and 48 hours of treatment. For phytohormone treatment, the roots were collected after 0, 1, 3, 6, 12, and 24 hours of treatment. The harvested samples were frozen immediately in liquid nitrogen and used for RNA isolation.

To analyze the root traits, seedlings were grown in rolled-up germination test paper in nutrient solution. After 15 days, root volume and root surface area were analyzed with WinRhizo Pro 2008a, an image analysis system (Regent Instr. Inc., Quebec, 13 Canada) with a professional scanner (Epson XL 1000; Japan). Root number, primary root length, root fresh weight, shoot fresh weight and shoot length (the length of the aerial part when the seedling was fully stretched) were measured manually. To analyze the effect of GA_1_ and GA_3_ on *ZmLBD33*-overexpressing plants, wild type (KN5585) and overexpression line 33-4 were germinated and grown in rolled-up germination test paper in nutrient solution containing GA_1_ or GA_3_ (0, 0.05, 0.1, 0.5, or 1.0 ng/mL). After 15 days, the shoot length, second leaf area (length × width × 0.75), root fresh weight, and shoot fresh weight were measured manually. Root traits were analyzed with WinRhizo Pro. For each treatment, at least 15 seedlings were used, and each treatment was repeated thrice.

For drought stress tests in individual pots, equal volumes of well-mixed soil, containing 200 mL of water, was put in each pot (length × width × height = 10 × 10 × 13 cm), and 5 seedlings of each line were grown in different pots. The control group was well watered, whereas the test group was not watered until it exhibited wilting. The shoot dry weight and root dry weight were measured. To analyze the soil moisture, 1 cm^3^ soil was sampled from each pot at three different stages (0, 10, and 15 days after planting).

For same-pot drought stress tests, larger pots (length × width × height = 54 × 28 × 4 cm) were used. Various transgenic plant lines and wild type plants were grown in the same pot. When test seedlings displayed significant wilting, photographs were taken and shoot dry weight was measured.

### Overexpression of *LBD5* and *LBD33*

The coding sequence of ZmLBD5 or ZmLBD33 was cloned into pCAMBIA3301 between BamH I and Sac I restriction sites and fused with HA and FLAG tags at N- and C- terminals, respectively, and expression was driven by the maize ubiquitin promoter. The vector was introduced into agrobacterium EHA105. Agrobacterium-mediated maize transformation was performed at Weimi Biotechnology (Jiangsu) Co. LTD, and the maize inbred line KN5585 was used as a recipient. Basta herbicide (0.3%, [v/v]) and PCR were used to identify transgenic plants.

### Yeast two-hybrid (Y2H) and bimolecular fluorescence complementation (BiFC)

To test whether ZmLBD5 and ZmLBD33 forms dimer with other LBDs or itself, the full codon regions of ZmLBD5, ZmLBD33, and ZmLBD44 were individually cloned into pGBK-T7 and/or pGAD-T7 vector and fused with BD and/or AD. BD-fused ZmLBD5, ZmLBD33, or ZmLBD44 was individually co-transformed with AD-fused ZmLBD5, ZmLBD33, and empty pGAD-T7 into the Y2HGold yeast strain. Yeast cells harboring pGBK-T7 and pGAD-T7 vectors were diluted to three concentrations and grown on nonselective (SD/-Trp/-Leu) or selective (SD/-Trp/-Leu/-His/-Ade) medium. A pGBKT7-53 and pGADT7-T combination was used as a positive control (+). A pGBKT7-Lam and pGADT7-T combination was used as a negative control (-). To test the effect of different regions, ZmLBD5 and ZmLBD33 were segmented into three parts (A: N-terminal C-block domain, B: GAS and leucine-zipper-like coiled-coil domain, and C: C-terminal domain) based on the genome wide analysis of LBD genes in maize (Zhang et al., 2014). For BiFC assays, the full codon regions of *ZmLBD5, ZmLBD33*, and *ZmLBD44* were individually cloned into pXYc104 and pXYn106 vectors and fused with the C-terminal (YC) and N-terminal (YN) of YFP. Then, indicated plastid combinations were co-transformed into agrobacterium cells. Transient expression was performed on *N. benthamiana* leaves. Primers and constructions are listed in Supplemental Table S4.

### Yeast one-hybrid (Y1H)

To investigate the binding of *ZmLBD5* and *ZmLBD33* to the promoters of candidate genes, *ZmLBD5, ZmLBD5AB, ZmLBD33*, and *ZmLBD33AB* were inserted into the pJG4-5 vector and fused with a TF-activating domain. The promoters (about 2,000-bp) of 15 candidate genes were inserted into pLacZi2u upstream of the lacZ reporter. Then, these vectors were co-transformed into the yeast strain EGY48, screened upon SD Base (without Trp and Ura, with glucose as the carbon source), and validated by PCR. The positive clones were then transferred to another SD Base [without Trp and Ura, replacing glucose with galactose and raffinose, and containing X-gal (5-bromo-4-chloro-3-indolyl-β-D-galactopyranoside)] for blue color development (Feng et al., 2016).

To investigate whether ZmLBD5 and ZmLBD33 have transcriptional inhibition activity, A, B, C, AB, BC, and full codon regions of *ZmLBD5* and *ZmLBD33* were amplified and fused with the *Gal4-AD* sequence. Then, the fused fragments were cloned into pGBKT7 and fused with the *Gal4-BD* sequence. *Gal4-AD* was cloned into pGBKT7 and fused with the *Gal4-BD* sequence, and complete *Gal4* was attained and used as a control. The transformation was conducted according to the manual of Yeast Protocols Handbook (Clontech). Primers and constructions are listed in Supplementary Table S4.

### Transient dual-luciferase expression assays

Reporters were constructed based on the pGreenII 0800-LUC vector and the effectors were constructed based on the pCAMBIA2300-eGFP vector. About 2,000-bp promoter fragments of candidate genes were amplified from B73 genomic DNA by PCR and cloned into pGreenII-LUC to drive the LUC reporter. *ZmLBD5* and *ZmLBD33* were cloned into pCAMBIA2300-eGFP and driven by the CaMV35S promoter. The empty vector pCAMBIA2300-eGFP was used as a control. Primers and constructions are listed in Supplementary Table S4.

Transient dual-luciferase assays were performed in tobacco leaves. A dual-luciferase assay kit (Vazyme, DL101-01) was used for enzymatic activity measurement. Three independent measurements were carried out for each analysis, and four biological repeats were performed.

### Measurement of stomatal density and stomatal aperture

At the three-leaf stage, the middle parts of the last fully expanded leaves were used for stomata measurement. The number of stomata in each microscopic field on abaxial leaf and adaxial leaf were calculated using an Olympus microscope (IX73, Japan) with a 10× objective lens. For each line, ten plants were selected for the counting and three replicates were performed. To investigate the ratio of open and closed stomata after dehydration, abaxial leaves were covered with clear nail polish 12 minutes after detachment. The shape of the stomata is stamped into the nail polish film during the rapid curing. The nail polish film was then stripped off and placed on a glass slide for microscopic observation with a 100× objective lens. For the assessment of stomatal closure dynamics induced by ABA and H_2_O_2_, the detached leaves were floated on stomatal opening buffer (10 mM Tris-HCl, pH 5.6, 10 mM KCl, and 50 μM CaCl_2_) for 3 h under light to induce the stomata to open to the maximum extent. Then, 10 μM ABA or 1 mM H_2_O_2_ was added to induce stomata closure and stomatal aperture was fixed at different time points using nail polish. Stomatal images were analyzed with ImageJ software to measure the aperture size. More than 30 stomata per sample were measured and each treatment included three replicates.

### RNA sequencing

RNA sequencing (RNA-seq) was performed on the transgenic lines 5-1, 33-5, and the wild type plants. Twelve-day-old seedlings were harvested for total RNA extraction. Sequencing and analysis were entrusted to Beijing Novogene company (https://www.novogene.com/) using the Illumina HiSeq 4000 sequencing platform.

### GA and ABA content

Endogenous GAs (GA_1_, GA_3_, GA_4_, and GA_7_) and ABA contents were measured in the overexpression lines 5-1 and 33-4, and in wild-type plants. Approximately 5 g of aerial tissue was harvested from 12-day-old seedlings grown under normal conditions (14-h light/10-h dark photoperiod at 28°C). Measurement of the GA and ABA content was entrusted to Convinced-Test (Nanjing) and analyzed by liquid chromatography-mass spectrometry (LC-MS). Two biological repeats were prepared, and three technical repetitions were performed for each sample.

### Real-time quantitative PCR

Total RNA was isolated with a Plant Total RNA Isolation kit (FOREGENE, RE-05014). Genomic DNA in samples was removed with RNase-free DNase I (Trans, GD201-01). RNA concentration was measured by spectrophotometer (NanoDrop 2000C). First-strand cDNA was synthesized using Prime Script RT reagent kit with gDNA Eraser (Takara, RR047A) from DNase I-treated RNA. Real-time quantitative PCR was performed using SYBR Green Fast qPCR Mix (ABclonal, RM21203) and a BioRad CFX96 machine. *ZmeF1α* was used as reference gene to normalize the expression of candidate genes. Primers are listed in Supplemental Table S5.

### Statistical Analysis

Unless noted otherwise, data are presented as the mean ± sd. Statistical significance was determined though one-way ANOVA analysis using GraphPad Prism (version 9.0). Variations were considered significant if P <0.05(*), 0.01(**) or 0.001(***).

### ACCESSION NUMBERS

Sequence data for genes and proteins presented in this article can be found in the *EnsemblPlants* database under the following accession numbers: *ZmLBD5* (Zm00001d029506), *ZmLBD33* (Zm00001d038717), *ZmLBD44* (Zm00001d023316), *ZmEF1a* (Zm00001d046449). The RNA-seq data was deposited in NCBI with BioProject ID: PRJNA715318.

## ACKNOWLEDGMENTS

We thank Dr. Peter Hedden from Rothamsted Research,UK for his inspiring work and the valuable suggestions on our research.

## Supplemental materials

**Figure S1.**
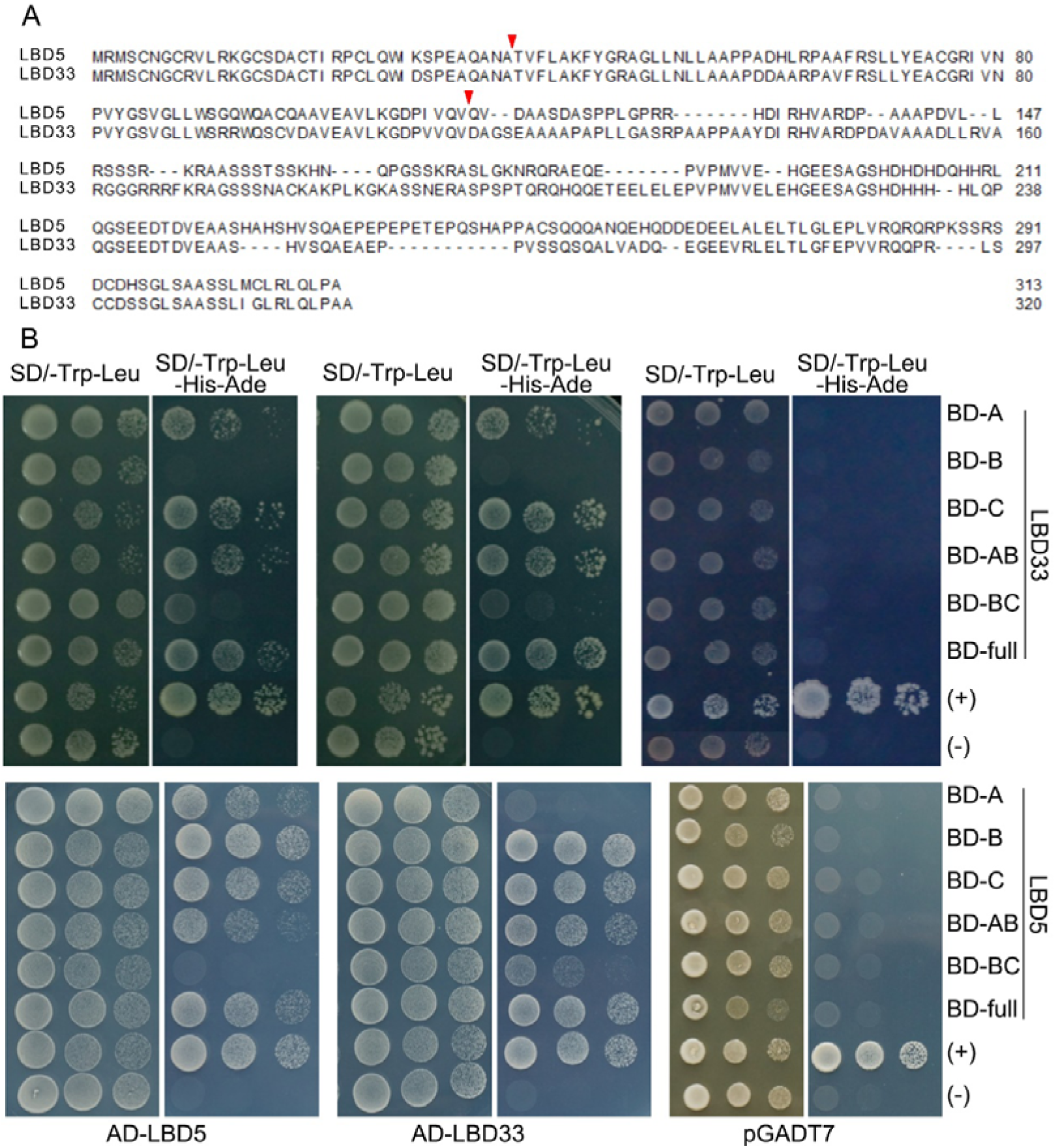
Investigation of the hetero- and homo- protein-protein interaction of ZmLBD5 and ZmLBD33. (A) Three segments of ZmLBD5 and ZmLBD33 were divided from red arrows indicated two sites. (B) Different segments of ZmLBD5 and ZmLBD33 were cloned into pGBK-T7 and fused with BD. Full codon region of ZmLBD5 and ZmLBD33 were cloned into pGAD-T7 and fused with AD. The yeast cells harboring the indicated plastid combinations were grown on nonselective (SD/-Trp/-Leu) or selective (SD/-Trp/-Leu/-His/-Ade) medium. Cells were diluted in three concentrations from left to right.

**Figure S2.**
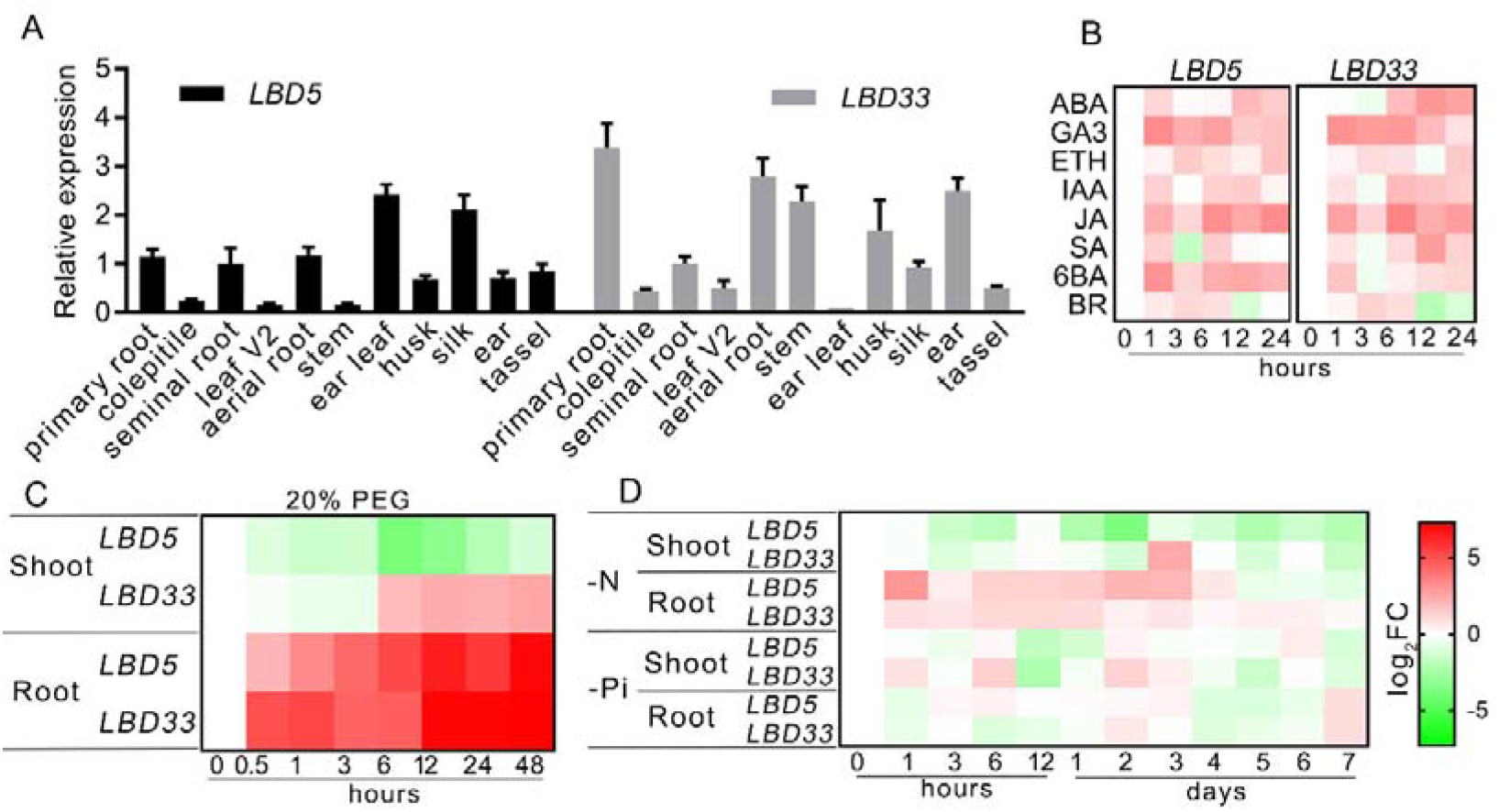
Expression of *ZmLBD5* and *ZmLBD33* upon different tissues and different stimuli. (A) Expression of *ZmLBD5* and *ZmLBD33* in different tissues, (B) upon different phytohormones, (C) upon PEG caused osmotic stress, (D) upon nitrogen or phosphorus deficiency. Fold change of expression relative to 0 point of treatment is converted to a logarithmic scale with base 2 and shown as different color in (B), (C), and (D).

**Figure S3.**
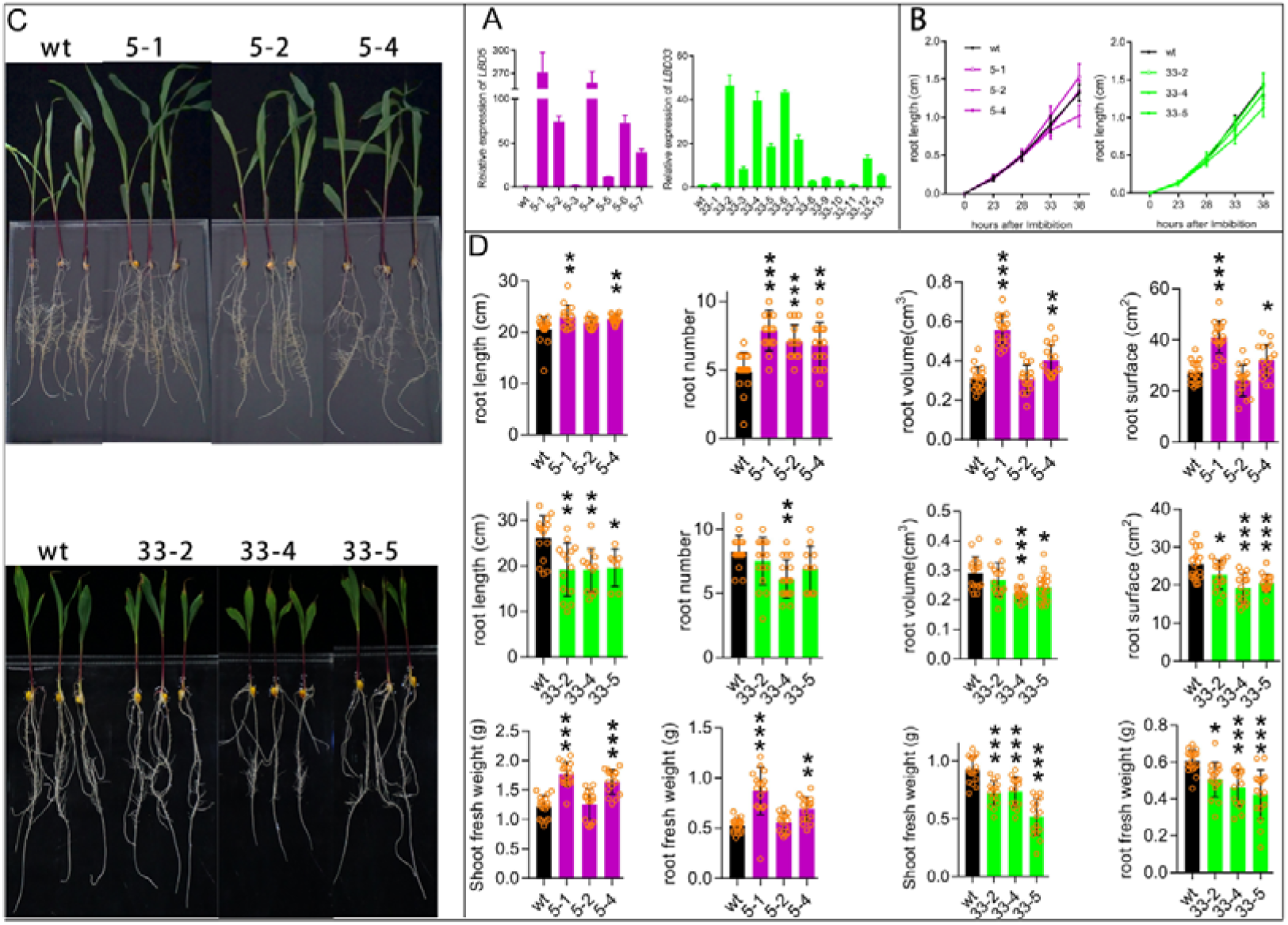
Phenotypic investigation of representative transgenic lines upon hydroponic conditions. (A) Relative expression of *LBD5* and *LBD33* in different transgenic lines. (B) The primary root length at different time after imbibition. (C) Phenotype and (D) quantitative description of primary root length, root number, root volume, root surface, shoot fresh weight and root fresh weight. Asterisks on bar represent the difference compared with wild type is significant.

**Figure S4.**
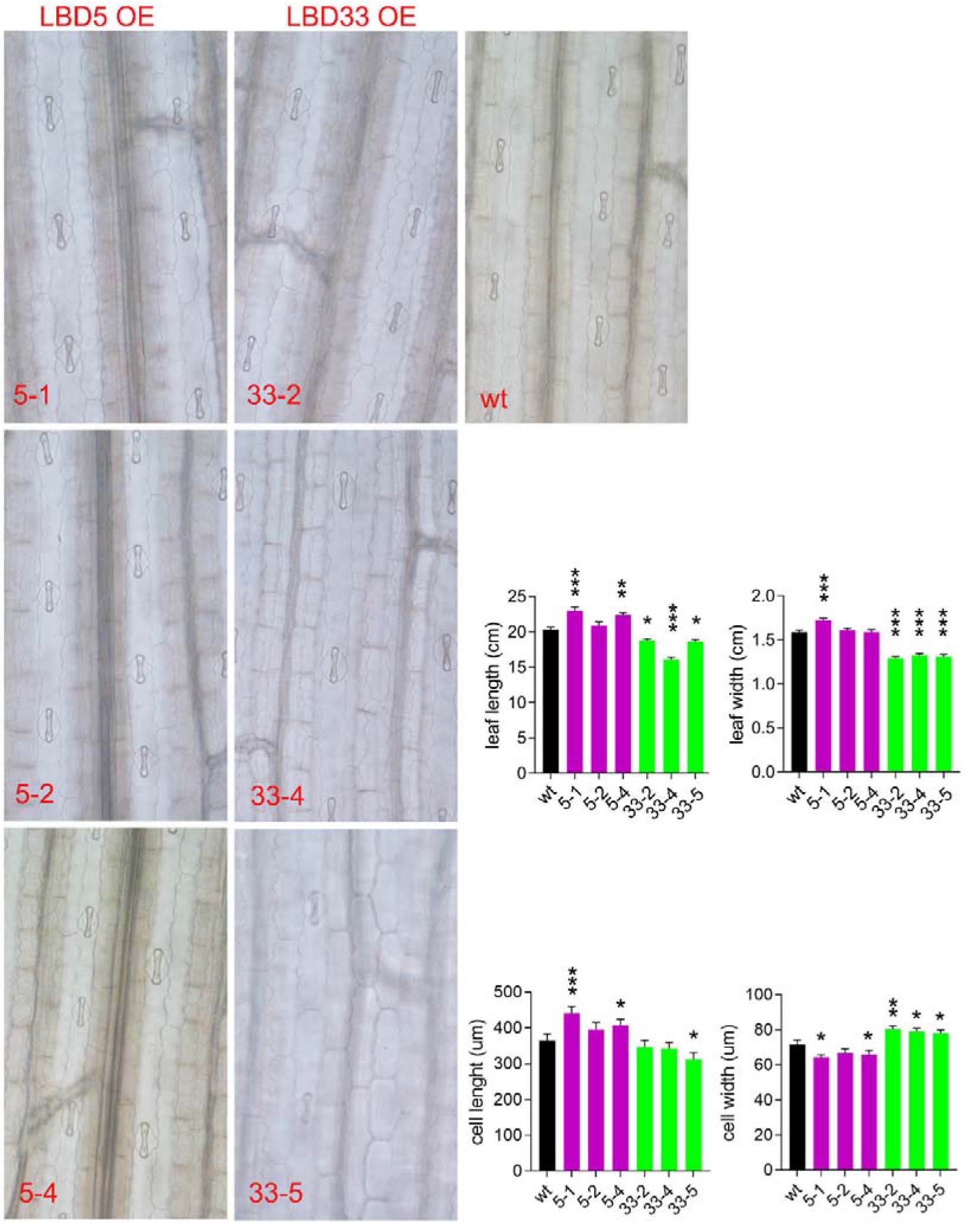
Microscale phenotype of third leaf on adaxial surface, cell length, cell width, leaf length and leaf width. Asterisks on bar represent the difference compared with wild type is significant.

**Figure S5.**
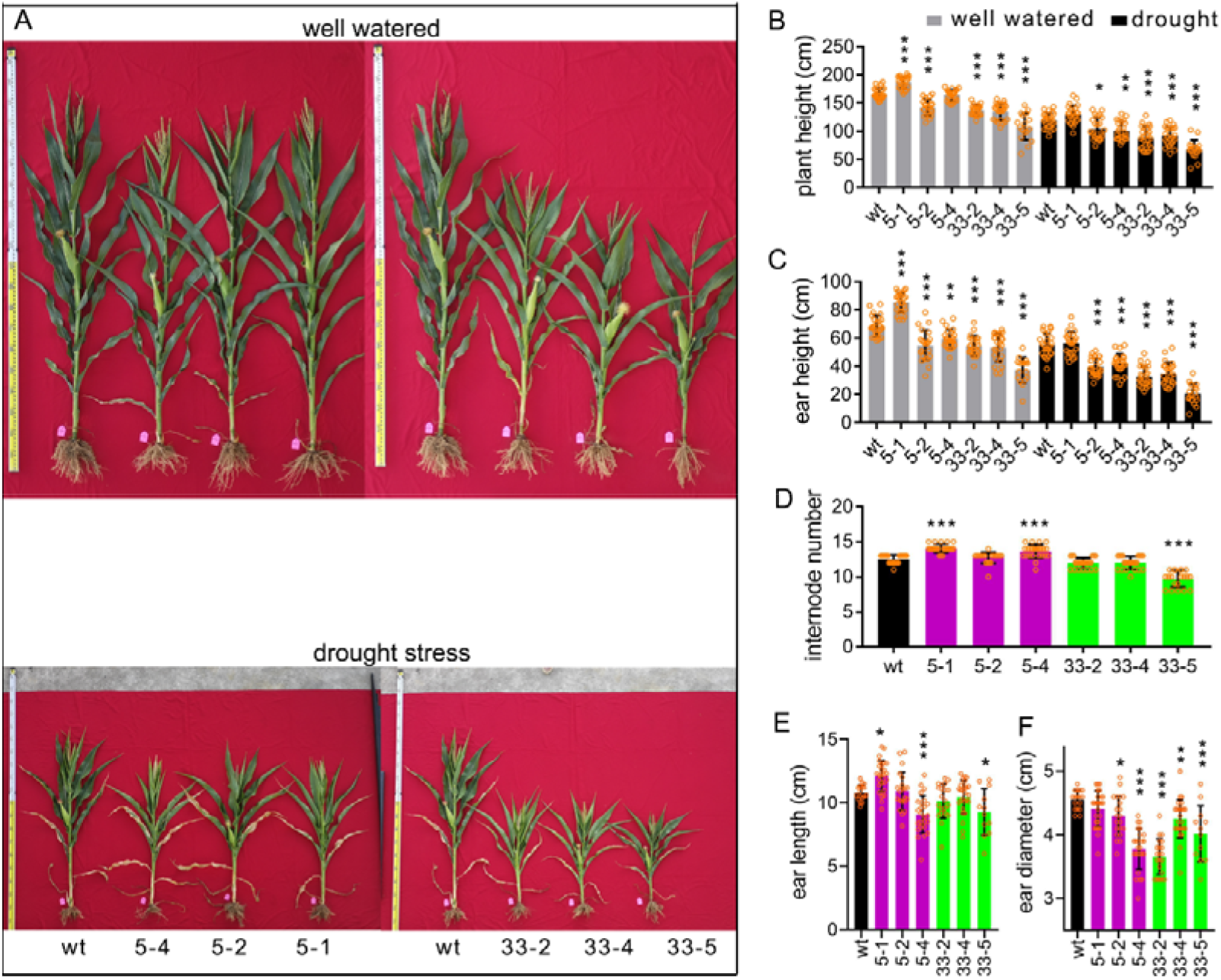
Phenotype of plants after tasseling. (A) Representative plants under well water condition. Plant height (B), Ear height (C), and internode number (D) of transgenic plants and wild type in field upon well water and drought conditions. Asterisks on bar represent the difference compared with wild type is significant.

**Figure S6.**
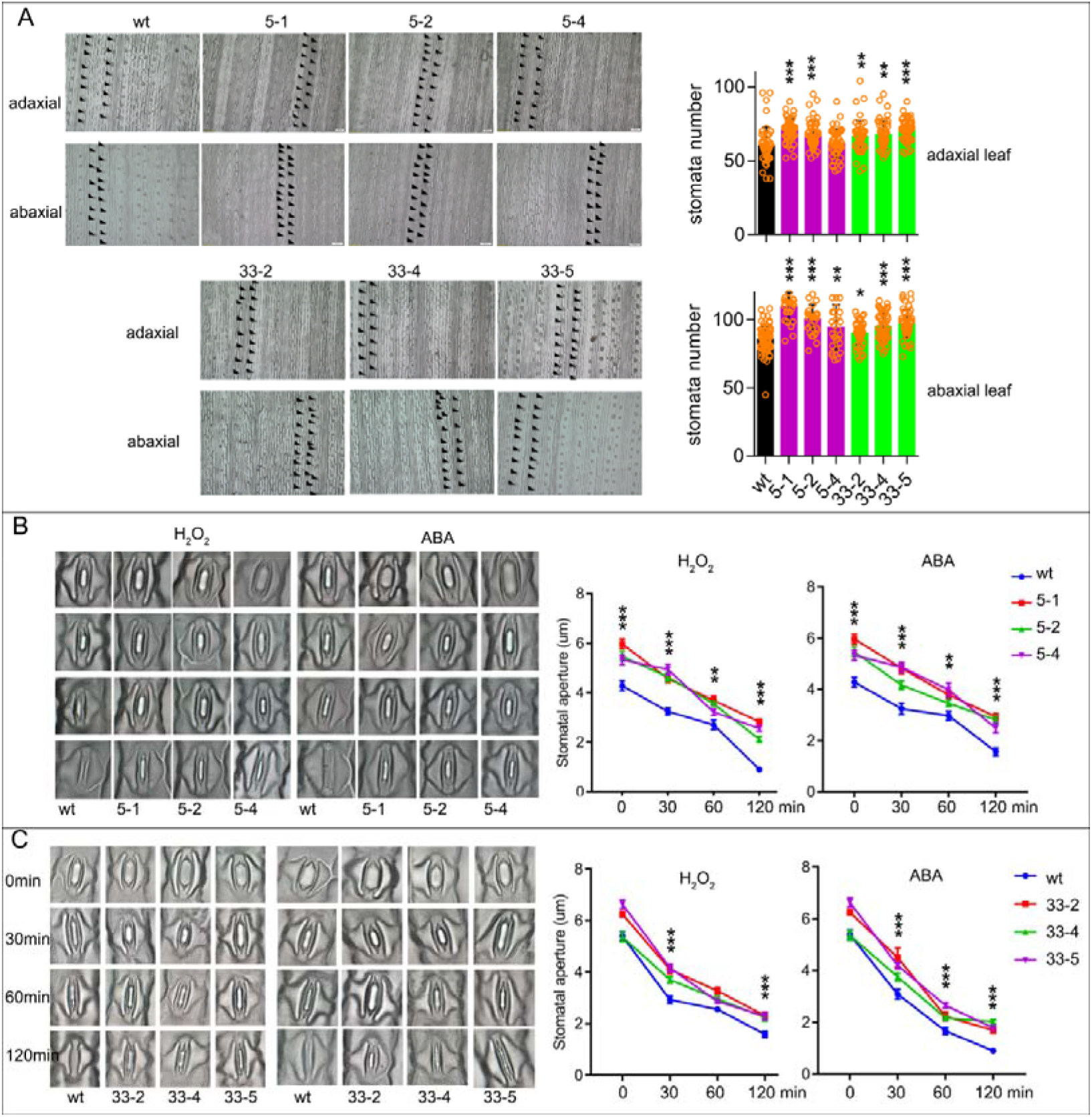
Stomata number and stomatal aperture upon ABA or H_2_O_2_ treatment. (A) Stomata number of the third leaf on abaxial surface and adaxial surface. Stomatal aperture of *LBD5* (B) and *LBD33* (C) overexpressed plants after ABA or H_2_O_2_ treatment. Asterisks on bar represent the difference compared with wild type is significant. In B and C, if all three lines were significant different with the wild type asterisk was labeled.

**Figure S7.**
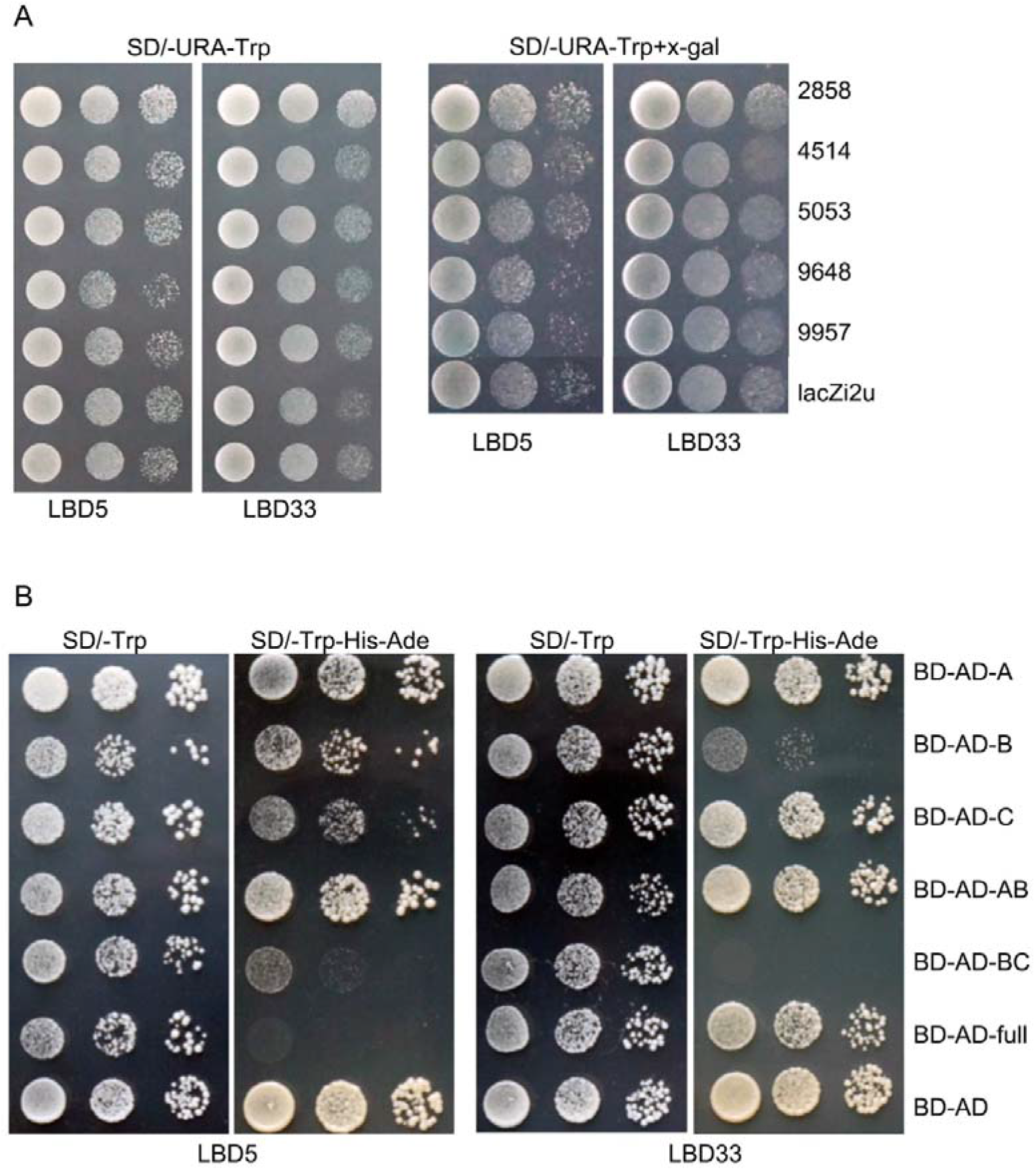
Yeast one-hybrid and potential transcriptional inhibition activity of LBD5 and LBD33. (A) The yeast cells harboring the indicated plastid combinations were grown on nonselective (SD/-Ura/-Trp) or color development (SD/-Ura/-Trp/+x-gal) medium. The last four number of the candidate gene name was used to represent corresponding gene. (B) Different segments of LBD5 and LBD33 were fused with Gal4-AD and cloned into pGBK-T7 to fuse with Gal4-BD. The yeast cells harboring indicated construct were grown on nonselective (SD/-Trp) or selective (SD/-Trp/-His/-Ade) medium. Cells were diluted in three concentrations from left to right.

**Table S1. internode length of plants after tasseling in field**.

**Table S2. DEGs in *LBD5-* and LBD33- overexpressing plant**.

**Table S3. Enriched GO terms and gene list in GGPP–CPP–kaurene/acid–GA metabolism and P450 members**.

**Table S4. Plasmid constructs and primers used in this study. Table S5. primers for real-time qPCR**.

